# Classical Mathematical Models for Prediction of Response to Chemotherapy and Immunotherapy

**DOI:** 10.1101/2021.10.23.465549

**Authors:** Narmin Ghaffari Laleh, Chiara Maria Lavinia Loeffler, Julia Grajek, Kateřina Staňková, Alexander T. Pearson, Hannah Sophie Muti, Christian Trautwein, Heiko Enderling, Jan Poleszczuk, Jakob Nikolas Kather

## Abstract

Classical mathematical models of tumor growth have shaped our understanding of cancer and have broad practical implications for treatment scheduling and dosage. However, even the simplest textbook models have been barely validated in real world-data of human patients. In this study, we fitted a range of differential equation models to tumor volume measurements of patients undergoing chemotherapy or cancer immunotherapy for solid tumors. We used a large dataset of 1472 patients with three or more measurements per target lesion, of which 652 patients had six or more data points. We show that the early treatment response shows only moderate correlation with the final treatment response, demonstrating the need for nuanced models. We then perform a head-to-head comparison of six classical models which are widely used in the field: the Exponential, Logistic, Classic Bertalanffy, General Bertalanffy, Classic Gompertz and General Gompertz model. Several models provide a good fit to tumor volume measurements, with the Gompertz model providing the best balance between goodness of fit and number of parameters. Similarly, when fitting to early treatment data, the general Bertalanffy and Gompertz models yield the lowest mean absolute error to forecasted data, indicating that these models could potentially be effective at predicting treatment outcome. In summary, we provide a quantitative benchmark for classical textbook models and state-of-the art models of human tumor growth. We publicly release an anonymized version of our original data, providing the first benchmark set of human tumor growth data for evaluation of mathematical models.

**Author Summary:** Mathematical oncology uses quantitative models for prediction of tumor growth and treatment response. The theoretical foundation of mathematical oncology is provided by six classical mathematical models: the Exponential, Logistic, Classic Bertalanffy, General Bertalanffy, Classic Gompertz and General Gompertz model. These models have been introduced decades ago, have been used in thousands of scientific articles and are part of textbooks and curricula in mathematical oncology. However, these models have not been systematically tested in clinical data from actual patients. In this study, we have collected quantitative tumor volume measurements from thousands of patients in five large clinical trials of cancer immunotherapy. We use this dataset to systematically investigate how accurately mathematical models can describe tumor growth, showing that there are pronounced differences between models. In addition, we show that two of these models can predict tumor response to immunotherapy and chemotherapy at later time points when trained on early tumor growth dynamics. Thus, our article closes a conceptual gap in the literature and at the same time provides a simple tool to predict response to chemotherapy and immunotherapy on the level of individual patients.

## Introduction

The growth of solid tumors and their response to therapy is hard to predict on the level of individual patients. Similar to other complex systems such as the climate [1,2] or stock markets [3], quantitative mathematical models can be used to describe and forecast the behavior of cancer: this is one of the main objectives of “mathematical oncology” [4,5]. Mathematical models of tumor growth kinetics have improved the understanding of underlying biological mechanisms. [6–8] In addition, they have resulted in a number of modeling approaches for cancer treatments including chemotherapy [9,10] and immunotherapy [11,12], improved drug dosage [13,14] and have yielded candidate biomarkers for treatment response [15]. The roots of tumor growth models go back to 1825, when Gompertz published a mathematical model to analyze the population growth [16]. He argued that the number of people alive as a function of their age *L*(*x*) declines faster than exponential functions which means that the death rate should be increasing with age. 135 years later, von Bertalanffy addressed the question of “why does an organism grow at all and why after a certain time, does its growth come to stop?” [17] By replacing its concept of an “organism” with a malignant tumor, the answer to this question resulted in a mathematical model for tumor growth. Tumor modeling provides information about the net tumor growth rate, facilitates their comparison among different tumor types [18] and makes it possible to predict the future growth of tumors [19].

A number of “textbook” models have been used in the past to approximate tumor growth with mathematical equations. In addition to the above-mentioned models by Gompertz and von Bertalanffy (each in a “classical” and a more general form), exponential and logistic models are standard approaches to describe tumor growth (Table 1) [20]. Exponential models are able to predict either exponential growth or decay depending on the absolute values of birth and death rates, and the resulting sign of (*birth rate* - *death rate*). Logistic models can simulate the fact that tumor growth is limited by nutritional, immunological or spatial constraints by including a carrying capacity into the model at which the tumor volume plateaus. This carrying capacity is included in the per capita growth rate, in line with the observation that tumor growth slows down when the tumor volume becomes large. [21] To be precise, the carrying capacity can be interpreted to comprise a number of biological constraints to tumor cell proliferation. These constraints include the availability of nutrients and oxygen and thus, the concept of tumor angiogenesis is implicit in the carrying capacity. In addition, the pressure of immune cells attacking tumor cells limits the niche the tumor cells can fill and thus, the concept of antitumor immune response is implicit in the carrying capacity. The Gompertz model is another model which illustrates the experimentally observed decrease in the growth speed of tumors. Similarly to the logistic model it has a sigmoid shape, hence representing limited tumor growth. Its main assumption is the exponential decay of the growth rate [18]. Since this model has been applied in many fields to various problems a few equivalent Gompertz models exist, differing in the chosen re-parametrization. Gompertz and von Bertalanffy growth models are two basic but important models which are commonly used to model tumor volume growth since they have outperformed exponential models in many cases in the past [22].

**Table 1.** Data Description. Five data sets were used in this study. The original number of patients in each data set and the treatment arm / subgroups are reported in this table. Two of the data sets have more than one treatment arm (Atezolizumab and Docetacxel) and the others have only one arm with a number of subgroups defined by clinical features. No. = Number, Pats. = Patients.

| Study ID | Cancer Type | Phase | No. Pats. | Treatment | Subgroup | No. Pats. per group |
| --- | --- | --- | --- | --- | --- | --- |
| NCT01846416<br>(GO28625)<br>**FIR** [52] | Non-Small Cell Lung Cancer | 2 | 138 | Atezolizumab* | MPDL3280A-1 | 31 |
|  |  |  |  |  | MPDL3280A-2 | 94 |
|  |  |  |  |  | MPDL3280A-3 | 13 |
| NCT01903993<br>(GO28753)<br>**POPLAR** [53] | Non-Small Cell Lung Cancer | 2 | 287 | Atezolizumab | - | 144 |
|  |  |  |  | Docetaxel |  | 143 |
| NCT02031458<br>(GO28754)<br>**BIRCH** [54] | Non-Small Cell Lung Cancer | 2 | 657 | Atezolizumab** | MPDL3280A-1a | 31 |
|  |  |  |  |  | MPDL3280A-2a | 79 |
|  |  |  |  |  | MPDL3280A-3a | 70 |
|  |  |  |  |  | MPDL3280A-1b | 104 |
|  |  |  |  |  | MPDL3280A-2b | 189 |
|  |  |  |  |  | MPDL3280A-3b | 184 |
| NCT02008227<br>(GO28915)<br>**OAK** [55] | Non-Small Cell Lung Cancer | 3 | 1182 | Atezolizumab | - | 609 |
|  |  |  |  | Docetaxel |  | 578 |
| NCT02951767<br>(GO29293)<br>**IMvigor 210** [56] | Bladder Cancer | 2 | 429 | Atezolizumab | - | 429 |

Unlike in other domains in which mathematical models and practice are strongly linked, the field of mathematical oncology is, by and large, somewhat disconnected from clinical practice of oncology. While in recent years, large quantitative data collections have deepened the genetic [23] and immunological [24–26] understanding of solid tumors, even well-established textbook models in mathematical oncology have not been linked with or validated in large amounts of quantitative real-world data. As a result, growth models that form a conceptual backbone of mathematical oncology have never been formally validated in large patient-derived datasets. In 2014, Benzekry *et al*. have systematically validated a range of textbook mathematical models on quantitative data obtained from two mouse models [20]. More recently, Vaghi *et al*. have extended that study and have validated classical growth models in 833 measurements in 94 animals [27]. These systematic large-scale approaches are highly important to link mathematical oncology to real-world data, but bear one major drawback: since almost all drugs that result in tumor control in mice fail in human experiments [28], mouse-based models are not suitable for human tumor growth estimations [29]. In addition, little validation of textbook models has been performed for tumors undergoing treatment, prompting caution whether unvalidated mathematical models have predictive power for clinical oncology.[30] While modeling of unabated tumor growth has academic relevance, fortunately untreated tumor growth for extended periods of time is rare in clinical practice [30]. Almost all patients with metastatic cancer undergo some type of systemic pharmacotherapy which slows down tumor progression [31].

In this study, we retrospectively collected quantitative measurements of tumor diameter changes over time from Non-Small Cell Lung Cancer (NSCLC) and bladder cancer patients from five large clinical trials. We systematically used this data with each of the standard mathematical models to address two questions: Firstly, how well can existing tumor growth models fit real-world data of patients undergoing treatment? (experiment #1) Secondly, how well can these models predict tumor growth at later disease stages when fitted to early-stage data? (experiment #2)

## Methods

### Ethics statement and data sharing

All experiments were conducted in accordance with the Declaration of Helsinki and the International Ethical Guidelines for Biomedical Research Involving Human Subjects by the Council for International Organizations of Medical Sciences (CIOMS). This study complies with the “Transparent reporting of a multivariable prediction model for individual prognosis or diagnosis” (TRIPOD) statement [32]. All data were obtained in an anonymized way through a proposal to F. Hoffmann-La Roche Ltd. through the platform “Clinical Study Data Request” (CSDR, www.ClinicalStudyDataRequest.com), which is now inactive and has been replaced by the Vivli platform (https://vivli.org, April 2021). Qualified researchers may request access to individual patient level data through the clinical study data request platform (https://vivli.org/). Further details on Roche’s criteria for eligible studies are available here (https://vivli.org/members/ourmembers/). For further details on Roche’s Global Policy on the Sharing of Clinical Information and how to request access to related clinical study documents, see (https://www.roche.com/research_and_development/who_we_are_how_we_work/clinical_trials/our_commitment_to_data_sharing.htm). The original proposal submitted to the CSDR platform is available in Annex 1. In order to enable reproduction of our experiments, we publicly release a fully anonymized subset of the data containing only the tumor volume measurements for the target lesion and the respective study and treatment arm (**Suppl. Table 1**).

### Data acquisition and preprocessing

We used data sets from five different clinical trials (**Table 1** and **Table 2**). The purpose of the original studies was the evaluation of the efficacy and safety of Atezolizumab (previously known as MPDL3280A), an immune checkpoint inhibitor directed against the Programmed Death Ligand 1 (PD-L1). In two out of the five trials (GO28753, GO28915), the performance of Atezolizumab was compared to Docetaxel, a chemotherapy drug. In the other three trials, all the participants received Atezolizumab as a treatment and the participants were further categorized into treatment arms or clinical subgroups as defined in the study protocols. One-dimensional longest diameter and shortest diameter of target and non-target lesions as manually measured on CT scans were available from the study database and were reported for each patient at different time intervals (**Figure 1A**). Because the shortest diameter was only available for a subset of patients, we used only the longest diameter (LD) and converted it to tumor volume (V) by *V* = *LD*^3^ * 0.5 as described before [33]. Using the maximum value of V in the whole data set, the volumes were normalized to be in the range of 0 and 1 for the whole dataset. Most patients in the data sets had multiple tumor lesions (primary tumor and/or metastases). For simplicity, we refer to these lesions as “tumors”. In the data set, one of these tumors for each patient was labeled as a “first target lesion” (‘INV-T001’), i.e. an easily measurable lesion for which the diameter was closely monitored over time. In addition, patients usually had one or more “non-target lesions”. In this study, we only used the target lesion which was labeled as for all the patients, as this was usually the tumor with the highest number of data points. Each patient has a different number of data points for this selected target lesion. While the intervals between data points were relatively similar (they were on average 50.62 days, with a standard deviation of 6.2 days), the absolute number varied (there were on average 3.63 data points for the selected target lesion per patient with a standard deviation of 3.22). Patients with few data points likely dropped out of the study early due to death or other reasons. To enable robust fitting of mathematical models to the data, we limited our data set to two sets of patients with three or more (or six or more, respectively) data points for the target lesion. Cumulatively, the original data sets had 2693 patients, of which 1472 had three or more data points and 652 had six or more data points available (**Figure 1B**).

**Figure 1.**
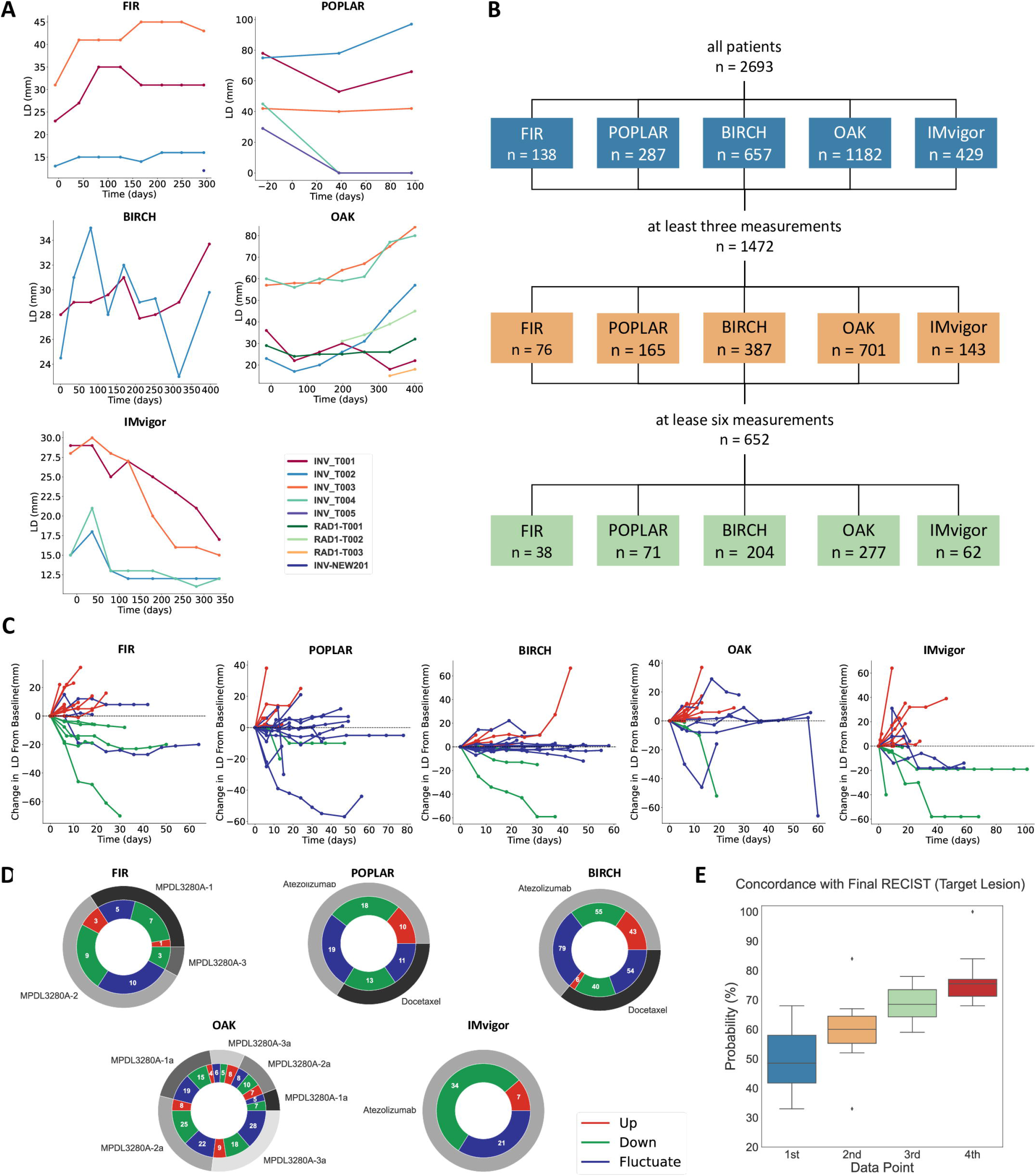
Data Description. (A) Longest tumor diameter over time for all lesions in representative patients in each data set. (B) Number of patients in each dataset. (C) Tumors can be categorized in three trajectory types based on their response to the treatment: Up, Down, Fluctuate. (D) Proportions of trajectory type in each dataset. (E) Initial RECIST status does not predict final RECIST status.

**Table 2.** Detailed summary of included studies. Data from five studies were used in this work. All studies can be identified either by their clinical trial registry number (“NCT…”) or by their Roche ID (“GO…”).

| Study | Design |
| --- | --- |
| NCT01846416<br>GO28625<br>FIR[52] | - Atezolizumab in PD-L1 + in NSCLC (n=138), Phase 2<br>- 1) patient with no first treatment<br>- 2) patients progress following platinum chemo<br>- 3) patients 2L + treated brain metastases<br>- ORR = 32 % /21% / 23% |
| NCT01903993<br>GO28753<br>POPLAR[53] | - After platinum failure: Atezolizumab or Docetaxel in NSCLC n=287; Phase 2<br>- 1) 144 in Atezolizumab group<br>- 2) 143 in docetaxel group<br>- OS 12.6 months / 9.7 months<br>- improvement in OS with higher PD-L1 expression<br>- Atezolizumab improved survival, correlated with expression PD-L1 |
| NCT02031458<br>GO28754<br>BIRCH[54] | - Atezolizumab in PD-L1 positive advanced or metastatic NSCLC n=667; Phase 2<br>- 1) 1L Atezolizumab<br>- 2) 2L Atezolizumab<br>- 3) 3L Atezolizumab<br>- ORR: 22% /19% / 18% |
| NCT02008227<br>GO28915<br>OAK[55] | - Atezolizumab vs Docetaxel advanced or metastatic NSCLC (2L) n = 1225, Phase 3<br>- OS better in Atezolizumab<br>- confirmed results of POPLAR study |
| NCT02951767<br>GO29293<br>IMvigor | - Atezolizumab in locally advanced or metastatic Bladder Cancer Phase 2<br>- 1) 1L atezolizumab<br>- 2) 2L atezolizumab after platinum based chemo<br>- study still ongoing (2020) |

### Patient categorization according to RECIST and trajectory type

For each patient, the ultimate response was encoded according to the response evaluation criteria in solid tumors (RECIST) system [34]. Based on the latest modification of this criteria (RECIST 1.1) [35], four tumor responses to treatments can be defined: Complete Response (CR, disappearance of all target lesions), Partial Response (PR, at least 30% decrease in sum of the longest diameters of target lesions in comparison to the baseline value), Progressive Disease (PD, at least an increase of 20% in the sum of the longest diameters of target lesions in comparison to baseline value) and Stable Disease (SD, when none of the above criteria fits to the tumor response). Because the RECIST system only assesses best response at discrete time points but does not categorize the full tumor volume trajectories, we additionally categorized the patients into three treatment response groups: “up”, “down” and “fluctuate” (**Figure 1C**). For this purpose, we calculated a vector containing the difference of each LD measurement at time point *t*+1 to its previous measurement at time point *t* for each patient. If the LD at *t* + 1 was bigger than at *t*, the difference would be positive and vice versa. Patients for whom only the shortest distance measurement was available were excluded from the analysis. The “up” category includes patients whose difference vector values are always positive and patients with a positive difference after the first measurement if the ratio between the sum of all positive values to the sum of all negative values is >2. The “down” category includes patients whose difference vector values are always negative or a negative difference after the first measurement if the overall ratio between the sum of all negative values to the sum of all positive values is >2. The “fluctuate” category contains all patients that correspond to neither up nor down categories. In all five studies, the pattern “up”, “down” and “fluctuate” was present (**Figure 1D**).

### Models and experimental design

We predefined six classical mathematical models to be fitted to the data (**Table 3**). Theoretically, there are two ways to integrate the effect of pharmacotherapy into models: either, explicit treatment-related arguments can be added to model equations, or, the effects of treatment can be implicit in the model. Here, we choose the implicit interpretation of treatment effects by assuming that therapies change either the growth or death rate of tumor cells or the carrying capacity of the tumor niche. For all models, the dependent variable is the volume of the tumor as a function of time. We subsequently performed to experiments: Experiment #1 was aimed at fitting models to the entire time series for each patient. The statistical endpoint for experiment #1 was the mean absolute error (MAE, also L1-norm, the lower the better, 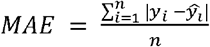, *y_i_* observed versus 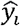 predicted values), the Akaike Information Criterion (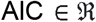, lower is better, 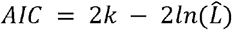, *k* is the number of parameters and 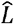 is the maximum value of the likelihood function for the model), Root Mean Square Error (RMSE, the lower the better, 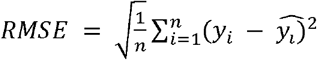 and R-squared fit (highest value is 1, higher is better, 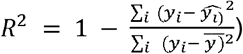. Experiment #1 was run on both patient sets separately: The set with all patients with three or more data points and the set with patients with six or more data points. Experiment #2 was aimed at fitting models to the early measurements for each patient, excluding the last three data points and subsequently estimating the predictive accuracy for the excluded data points. The statistical endpoint for experiment #2 was the MAE. Experiment #2 was only run on the set of patients with six or more data points.

**Table 3.** Model Description and interpretation of the parameters. For all differential equation models in the current study, the model name, equations and variables are listed. *birth rate and growth rate can be combined to one parameter, the effective growth rate.

| Model name | Solution of the differential equation | Differential equation (with initial condition $V(0) = V_0$) | Parameter description | Ref. |
| --- | --- | --- | --- | --- |
| Exponential | $V(t) = N_0 e^{(\alpha - \beta)t}$ | $\frac{dV}{dt} = (\alpha - \beta) V$ | $\alpha$ [$time^{-1}$]: birth rate*<br>$\beta$ [$time^{-1}$]: death rate* | [41] |
| Logistic (Verhulst) | $V(t) = \frac{V_0 K e^{\gamma t}}{K - V_0(1 - e^{\gamma t})}$ | $\frac{dV}{dt} = \gamma N (1 - \frac{V}{K})$ | $\gamma$ [$time^{-1}$]: max. net growth rate<br>$K$ [$mm^3$]: carrying capacity | [41] |
| Gompertz | $V(t) = \exp(\frac{\delta}{\gamma} + (\ln(V) - \frac{\delta}{\gamma})e^{-\gamma t})$ | $\frac{dV}{dt} = V(\delta - \gamma \ln V)$ | $\gamma$ [$mm^3^{-1} time^{-1}$]: max. net growth rate<br>$\delta$ [$time^{-1}$]: constant | [19,41, 57] |
| General Gompertz | | $\frac{dV}{dt} = V^\lambda (\delta - \gamma \ln V)$ | $\gamma$ [$mm^3^{-1} time^{-1}$]: max. net growth rate<br>$\delta$ [$time^{-1}$]: constant<br>$\lambda$ : constant | [41] |
| Classic von Bertalanffy | $V(t) = \left( \frac{\alpha}{\beta} + \left( V_0^{\frac{1}{3}} - \frac{\alpha}{\beta} \right) e^{-\frac{1}{3}\beta t} \right)^3$ | $\frac{dV}{dt} = \alpha V^{\frac{2}{3}} - \beta V$ | $\alpha$ [$time^{-1}$]: birth rate<br>$\beta$ [$time^{-1}$]: death rate | [41] |
| General von Bertalanffy | $V(t) = \left( \frac{\alpha}{\beta} + \left( V_0^{1-\lambda} - \frac{\alpha}{\beta} \right) e^{-\beta(1-\lambda)t} \right)^{\frac{1}{1-\lambda}}$ | $\frac{dV}{dt} = \alpha V^\lambda - \beta V$ | $\alpha$ [$time^{-1}$]: birth rate<br>$\beta$ [$time^{-1}$]: death rate<br>$\lambda$ : constant | [41] |

### Fitting and implementation

All model fitting procedures were implemented in Python 3.7. In particular, we used differential evolution to generate the initial data points for the differential equations. Differential evolution is a stochastic population based method which is used for global optimization problems [36]. Based on these initial guessed parameter values, the “Curve_fit” function (from the python package “scipy”) is used to fit the model parameters to the experimental data. This function uses the Trust Region Reflective (trf) algorithm with the non-linear least squares loss function to find an optimal fit of the model parameters to the data points. The inputs to this function are the sorted time and its corresponding tumor volume measurements, the respective mathematical function to fit the data and the maximum number of iterations (we used 1000 iterations in this study). The output of the “Curve_fit” function is the calculated optimum parameters for the selected mathematical function. Having these parameters, it is possible to predict the volume values for each time point and then evaluate the goodness of the fit. The source codes are publicly available at https://github.com/KatherLab/ImmunotherapyModels.git

## Results

### Early RECIST status does not correspond well with ultimate treatment response

In clinical routine and clinical trials, RECIST response at early time points during treatment is often used to determine whether a given treatment should be continued [34]. If these initial RECIST results perfectly matched the ultimate RECIST, there would not be a need for more mathematical prediction models. Therefore, we systematically compared the RECIST status at the first, second, third and fourth tumor size evaluation for each patient with the “final” RECIST status as defined in the study protocol. In all treatment arms, we found an imperfect overlap between early and final measurements. Overall, the median concordance between first, second, third and fourth data point and final RECIST was 53.5, 64.0, 63.5 and 78.0, respectively. The same pattern was seen for the concordance between early and final RECIST calculated for only one target lesion (**Figure 1E**). Hence, the RECIST classification can be a useful tool to assess therapy response status, but it might be insufficient for therapy response estimation at an early therapy stage. These findings provide a rationale for the use of mathematical models to improve response prediction. In addition, we compared statistically the correlation between the RECIST standard classification categories (CR/PR, SD and PD) with the developed grouping methods (up, down and fluctuate). As the results are summarized in **Suppl. Figure 1** both grouping systems are partially correlated (PD is mostly overlapping with “up”, PR/CR with “down” and SD with “fluctuate”). However, the correlation was not perfect and particularly in the OAK study, 37 patients from 95 down category patients are classified as PD and 87 patients out of 133 patients in fluctuate category are classified as PD. This comparison shows that while RECIST is the standard classification system in clinical routine, our grouping method does provide an additional perspective on tumor response categories.

### The Gompertz model outperforms other models when fitting clinical data points

We tested how well classical differential equation models (**Table 3**) can fit tumor volume trajectories under immunotherapy and chemotherapy. To compare these models, we first fitted them to all available data points for all patients with at least six measurements (experiment #1). This set the stage for experiment #2 (**Figure 2A**), in which models were fit to all points except the three last points and the predictive power was assessed for each model. In experiment #1, we found that all models provided a good fit to most data points, but the number of poorly fitted points differed between the models. Overall, the General Bertalanffy, the Gompertz and the General Gompertz model had the lowest number of poorly fitted data points (**Figure 2C**). We quantified this by calculating multiple metrics for the goodness of fit for each model, for each study arm, further stratifying patients in each study arm by the ultimate RECIST response. Again, we found that the General Bertalanffy, the Gompertz and the General Gompertz model consistently outperformed more simple models. (**Figure 3A and B**) The exponential model yielded the worst fit for 13 out of 19 patient groups in this analysis (**Figure 3A**). To rule out a selection bias, we repeated experiment #1 with all patients with at least three measurements, yielding comparable results (**Suppl. Figure 2A and B**). Due to their higher degree of freedom, complex models always yield a better fit than simple models to any data set. To account for this, we assessed the Akaike Information Criterion (AIC) which incentivises goodness of fit but penalizes model complexity. We found that according to the AIC, the General Bertalanffy model consistently yielded the poorest performance compared to the other models (**Figure 3C and D**). This observation also held when all patients with three or more measurements were considered (**Suppl. Figure 2C and D**). However, the Gompertz model had a low (good) AIC for most study arms, showing that this model give a good balance between goodness of fit and model complexity. To rule out that these effects were obtained by sub-stratifying patients according to their final RECIST status, we repeated experiment #1 with patients sub-stratified as “up”, “down” and “fluctuate”, thereby considering the shape of the whole timeline for each patient. Again, we found that the General Bertalanffy, the Gompertz and the General Gompertz model consistently outperformed the exponential model, the logistic model and the Classic Bertalanffy model in terms of Mean Absolute Error (**Suppl. Figure 3A and B**), the Root Mean Square Error (**Suppl. Figure 3C and D**) and the R-squared Error (**Suppl. Figure 3E and F**). In particular, this was the case for “fluctuating” patients which for the most clinically interesting group of patients (**Suppl. Figure 4**). For the “up” and “down” patient groups, the fitted model parameters were generally in a close range. For the “fluctuating” patient group, the fitted model parameters showed a higher variability between the patients, indicating the difficulty of to fit these trajectories (**Suppl. Table 2)**. When penalizing for model complexity by using the Akaike Information Criterion, again the Gompertz model provided the best balance between goodness of fit and model complexity (**Suppl. Figure 3 G and H**). In summary, the Gompertz model adequately fitted the response to immunotherapy and chemotherapy across a range of clinically relevant populations, while having only two free parameters (**Table 3**).

**Figure 2.**
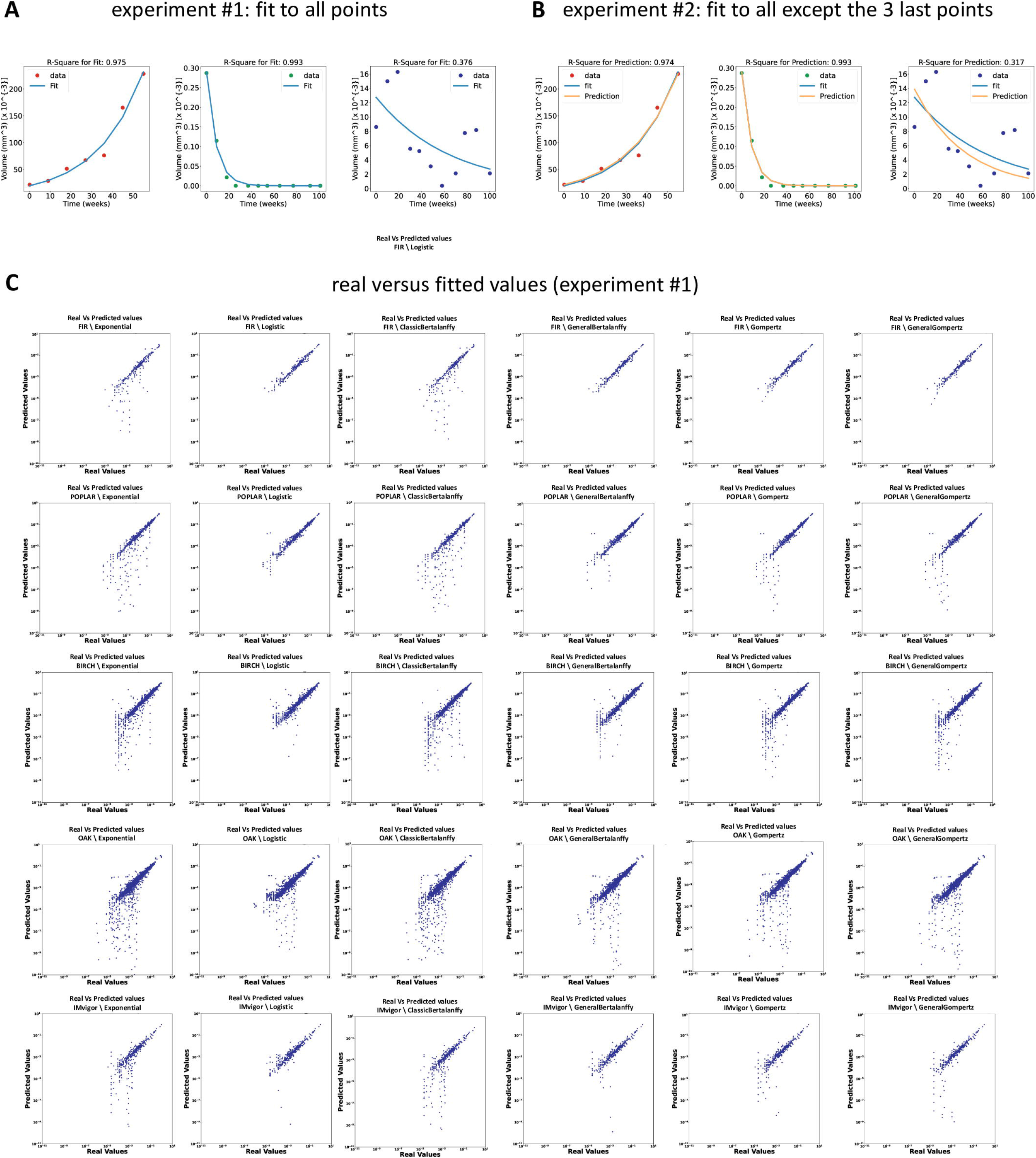
Experimental design and model fit. (A) In experiment #1, models were fitted to all available data points for each patient (only for patients with at least 3 or 6 data points, respectively). In experiment #2, models were fitted to all but the last 3 data points for all patients with at least 6 data points. Then, the predictions for the last 3 data points were compared with the actual values. (B) Fit and prediction for three representative patients. (C) Plot of real data points and fitted data points for all models for all studies. A larger deviation from the diagonal indicates a worse fit. Models with a “raincloud” appearance systematically underestimate true tumor volume.

**Figure 3.**
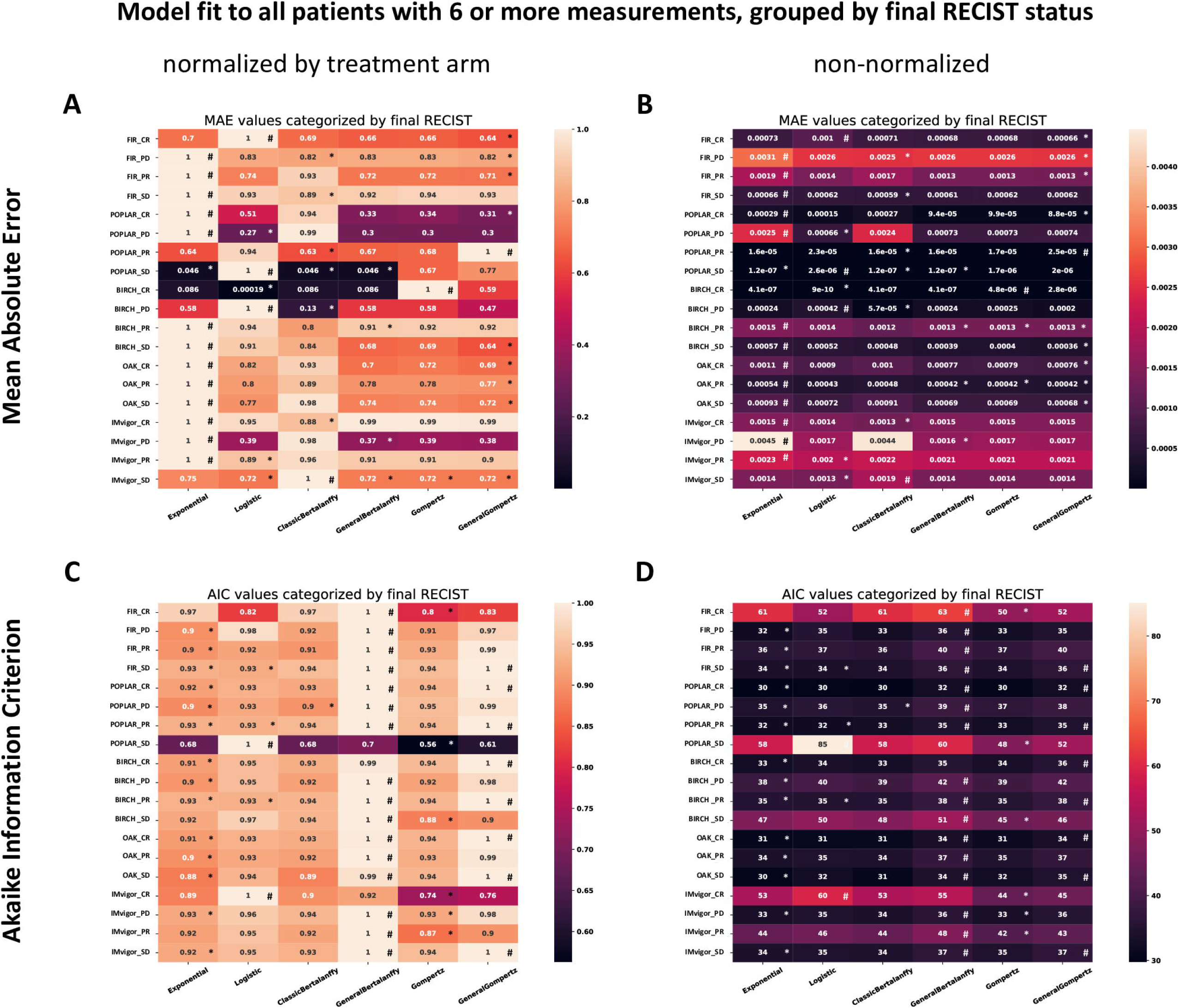
Head-to-head comparison of all models. (A) Model fit for all treatment arms in all trials, stratified by final RECIST, for all models. The loss function is the Mean Absolute Error (MAE, L1-Loss), after row-wise normalization. (B) Corresponding plot without row-wise normalization, showing the raw MAE. The worst MAE in each figure is indicated with “#” and best one is indicated with “*”. (C) Corresponding plot showing the Akaike Information Criterion (AIC) which penalizes models with a large number of free parameters, row-wise normalized. (D) Corresponding plot without row-wise normalization.

### Differential equation models can predict tumor response from early time points

While it is important to assess a model’s ability to fit a tumor volume timeline a posteriori, a more clinically relevant problem is to predict final treatment response based on early tumor behavior under therapy. Therefore, we investigated if these models can predict the last data points when only fitted to early treatment response. To investigate this, we held out the three last data points on any given patient, fit the model to all remaining (early) data points and evaluated the mean absolute error from extrapolation to the holdout test measurements (experiment #2). Interestingly, we found that in most patient groups in most treatment arms the holdout data points could be very well predicted with this approach. A remarkable exception was the Classic Bertalanffy Model, which yielded the worst fit on the last three points as assessed by the Mean Absolute Error (**Suppl. Figure 5A and B**). Overall, the best models for predicting holdout measurements were the General Bertalanffy and the Gompertz model (**Suppl. Figure 5A and B**). When analyzing the predictions of the exponential model (**Figure 4A**) and the General Bertalanffy model (**Figure 4B**) in more detail, we found that for the “up” and “down” patients, the exponential and the General Bertalanffy model visually recapitulated the trajectory of the tumor volume. A notable exception are U-shaped curves present in some of the “fluctuating” patients (**Figure 4A and B**).

**Figure 4.**
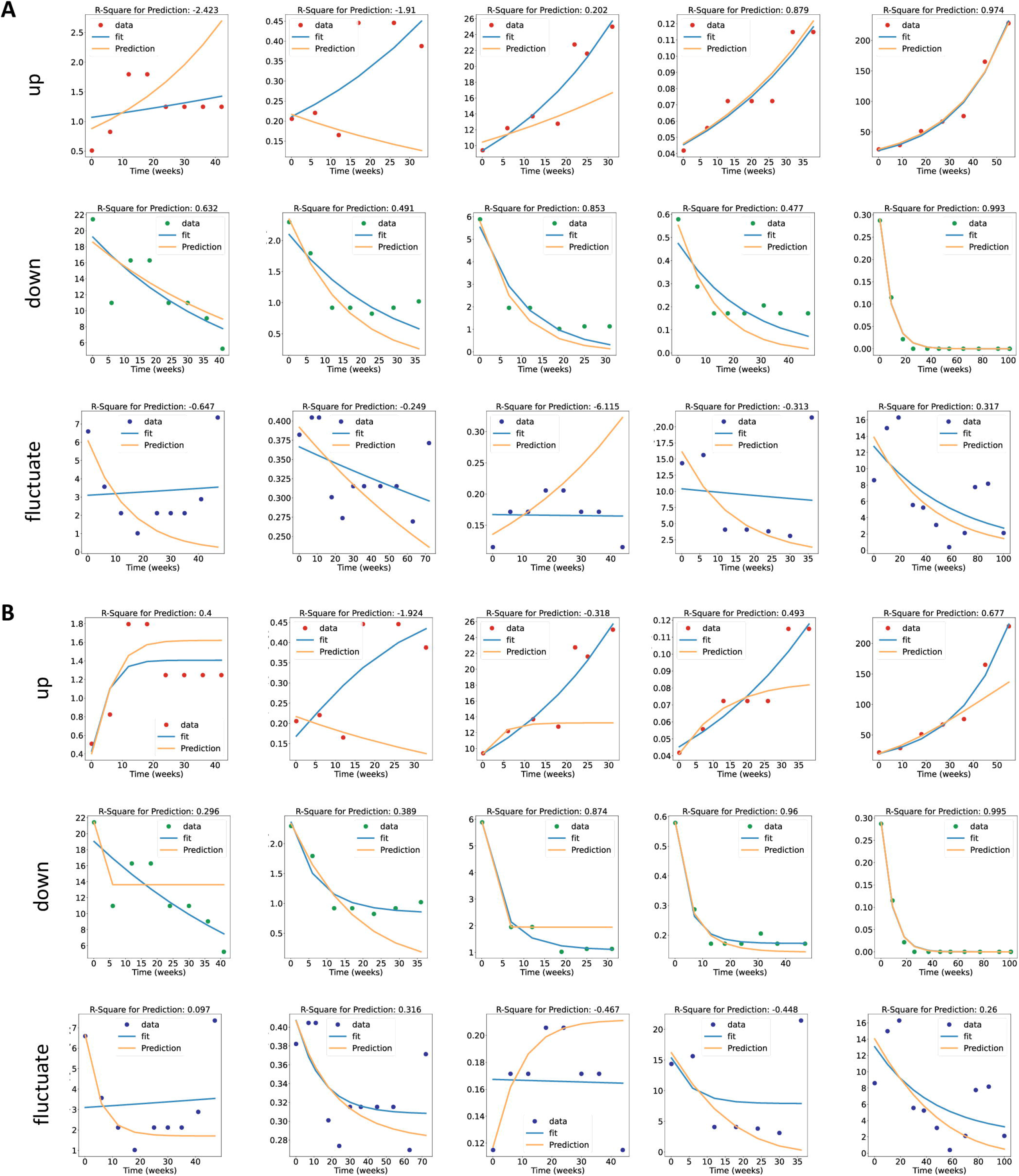
Fit of the exponential model and the General Bertalanffy model to unseen data. (A) Fit (blue) of the exponential model to the full timeline of representative patients with “up”, “down” and “fluctuate” trajectories. For the same patients, the prediction (yellow) is shown which was fitted to all points except the last three data points. (B) Corresponding plot for the General Bertalanffy model. The y axis is the relative tumor volume with respect to the largest tumor in the whole dataset, shown as 10^-3.

## Discussion

Cancer immunotherapy with immune checkpoint inhibitors is now an established part of the therapeutic arsenal for solid tumors [37]. Patterns of response to this class of drugs are more complex than for classical chemotherapy [38]. Previous studies have at length discussed new response trajectories such as hyperprogression, pseudoprogression [38,39] or delayed response [40] in immune checkpoint inhibitors. Accordingly, simple assessment systems for treatment response such as RECIST are not ideally suited to predict future treatment response for a given patient. Although mathematical models of tumor growth have been used for decades to understand mechanisms of tumor progression and treatment response, they have not been systematically validated in human real-world data of patients undergoing systemic treatment. To our knowledge, the only large systematic evaluation of these models have been performed on mouse tumors [20,27], which function merely as a proxy for human tumors. Moreover, although immunotherapy is a cornerstone of cancer treatment and classical mathematical models are in principle useful to model cancer growth under therapy, they have not been previously applied to large cohorts of patients under immunotherapy. In this study, we present a systematic application of mathematical tumor growth models on a large human dataset of patients undergoing immunotherapy and chemotherapy. We restricted our analysis to six consensus mathematical models selected from [41]. We show that in particular the Gompertz model and the General Bertalanffy can successfully fit the tumor growth trajectory and provide an accurate prediction of ultimate treatment response on the basis of early treatment data. However, we also show that the fit for “fluctuating” patients is lower in all models, and fully U-shaped tumor growth trajectories could not be fitted at all. Comparison of the results between experiment #1 and #2 shows that models perform better if all the data points are used. However, from the clinical point of view, it is very useful if a model can predict the final response points from the early treatment response. This highlights the usefulness of stratifying patients into different categories and, in the future, of using more sophisticated models which can overcome this limitation. Our findings mirror a previous study by Benzekry et al. who demonstrated that the Gompertz model provides a good approximation of tumor growth in mice. [20] Therefore, our study provides a potential bridge between textbook models of mathematical oncology and oncology practice today, providing evidence that simple mathematical functions can be used to predict immunotherapy response in most patient subsets.

A structural limitation to our study lies in the circumstance that the simple modelling of tumor growth or decay might not be the best predictor for the overall therapy outcome. Although the assessment of tumor growth might be useful to evaluate the drug or therapy regime response, it does not provide overall survival prediction for individual patients. Tumors might show a positive therapy response, but at the same time patients might die from adverse therapy events, infections or other therapy-related problems. Consequently, mathematical models which are solely based on tumor growth data should only be used together with other prognostic and predictive factors in clinical routine. Another limitation is the fact that by setting a threshold of at least six measurements at six points of time per patient, we had to exclude a part of patients from our final analysis. We mitigated this problem by repeating the analysis for patients with at least three data points, but this could still represent a selection bias by neglecting early study drop-outs and early cancer-related deaths. Other data-related limitations are that for some patients, only very few points can be present during the initial dynamics which might create problems. In the future, the availability of more complex datasets could allow researchers to build more complex models, thereby capturing more nuanced details of tumor growth. In practice, this is limited by the availability of structured data in oncology. In addition, in line with previous studies performed on mouse data, we used very simple mathematical models in this study [20,27]. Such models are a strong simplification of the reality of solid tumors, which are multicellular structures with a distinct spatial architecture [42]. Fundamentally, the key question is: how granular should a model be? This has been discussed extensively in the literature [8,43–47]. More complex models have been proposed for modeling tumor growth under immunotherapy which could improve the fit to the data, for example the Kuznetsov model [12] and game theoretical models [48–50]. As a starting point for the analysis of more complex models of computational oncology in real-life human datasets of various cancer types, we provide our raw data for re-use by other groups. In addition to non-spatial models like the ordinary differential equation (ODE) models in this study, other studies have explored the use of spatial models in the context of cancer immunotherapy. [45,46] However, in these studies we found that it is very hard to fit the parameters of spatial models to clinical routine data. Even simple spatial models have >25 free parameters, which means that for every patient at least 25 measurements are needed (ideally much more). In comparison, the ordinary differential equation (ODE) models in our study are much simpler and they only have two or three free parameters. This simplicity enables fitting the model parameters to routine clinical data such as the databases used in our study. Furthermore, the use of non-spatial models is supported by theoretical considerations. Solid tumors consist of billions of cells which show some mobility in the immediates spatial vicinity. Tumors are not perfectly homogeneous in the spatial dimension, but if we assume that the relevant biological processes are sufficiently similar in distinct parts of the tumor, spatial patterns do not have to be explicitly modeled, but can be implicit as in ODE models. Ultimately, complex spatial models and simplistic ODE models are both very valuable tools which could be implemented in the clinic in different situations. Our present study provides the first large-scale evidence for the usefulness of ODE models. Future studies should investigate more complex models in similar experimental approaches. In general, clinical utility remains the ultimate benchmark, as was pointed out by Gerlee [51], “a model that is disconnected from reality in terms of mechanisms and dynamics is acceptable, as long as it does the job of predicting”.

Ultimately, after refinement and prospective validation, such models could conceivably be used in the clinic to provide guidance on treatment recommendations for cancer patients. Unlike molecular biology-based biomarkers in the field of oncology, mathematical models could potentially improve response prediction for individual cancer patients based on ubiquitously available routine data.Tables

## Supporting information

Inventory of Suppl. Material

## Additional information

## Acknowledgements

The authors thank F. Hoffmann-La Roche Ltd. for sharing the raw data through the platform “Clinical Study Data Request” (CSDR, www.ClinicalStudyDataRequest.com).

## Legend of supporting information

**Suppl. Figure 1: Statistical comparison between the “up”/”down”/”fluctuate” and the standard RECIST-based grouping “CR/PR”/”CR”/”PD”.**

**Suppl. Figure 2: Model fit to all patients with three or more measurements.** (A) Model fit for all treatment arms in all trials, stratified by final RECIST, for all models. The loss function is the Mean Absolute Error (MAE, L1-Loss), after row-wise normalization. (B) Corresponding plot without rowwise normalization, showing the raw MAE. The worst MAE in each figure is indicated with “#” and best one is indicated with “*”. (C) Corresponding plot showing the Akaike Information Criterion (AIC) which penalizes models with a large number of free parameters, row-wise normalized. (D) Corresponding plot without row-wise normalization.

**Suppl. Figure 3: Model fit to all patients grouped by trajectory type and additional loss functions.**

**Suppl. Figure 4: Goodness of fit for all models, all trial arms, all patient groups.**

**Suppl. Figure 5: Goodness of fit for unseen data points for each model.** Results of experiment #2.

**Annex 1: Original data sharing request**

**Suppl. Table 1: fully anonymized subset of the data containing the tumor volume measurements for the target lesion and the respective study and treatment arm.**

**Suppl. Table 2: Distribution of the parameters for different types of trajectories in all the 5 datasets calculated by the examined 6 mathematical models.** Table 3 is a reference to the used parameters for each function.

## References

1. Manabe S, Stouffer RJ. A CO2-climate sensitivity study with a mathematical model of the global climate. Nature. 1979;282: 491–493.

2. Flores JC, Bologna M, Urzagasti D. A mathematical model for the Andean Tiwanaku civilization collapse: climate variations. J Theor Biol. 2011;291: 29–32.

3. Ledoit O, Santa-Clara P, Wolf MN. Flexible Multivariate GARCH Modeling with an Application to International Stock Markets. SSRN Electronic Journal. doi:10.2139/ssrn.311514

4. Rockne RC, Scott JG. Introduction to Mathematical Oncology. JCO Clin Cancer Inform. 2019;3: 1–4.

5. Anderson ARA, Quaranta V. Integrative mathematical oncology. Nat Rev Cancer. 2008;8: 227–234.

6. Araujo RP, McElwain DLS. A history of the study of solid tumour growth: the contribution of mathematical modelling. Bull Math Biol. 2004;66: 1039–1091.

7. Araujo A, Cook LM, Lynch CC, Basanta D. Size Matters: Metastatic Cluster Size and Stromal Recruitment in the Establishment of Successful Prostate Cancer to Bone Metastases. Bull Math Biol. 2018;80: 1046–1058.

8. Enderling H, Wolkenhauer O. Are all models wrong? Comput Syst Oncol. 2020;1. doi:10.1002/cso2.1008

9. Vainstein V, Kirnasovsky OU, Kogan Y, Agur Z. Strategies for cancer stem cell elimination: insights from mathematical modeling. J Theor Biol. 2012;298: 32–41.

10. Powathil GG, Gordon KE, Hill LA, Chaplain MAJ. Modelling the effects of cell-cycle heterogeneity on the response of a solid tumour to chemotherapy: biological insights from a hybrid multiscale cellular automaton model. J Theor Biol. 2012;308: 1–19.

11. Kogan Y, Halevi-Tobias K, Elishmereni M, Vuk-Pavloviċ S, Agur Z. Reconsidering the paradigm of cancer immunotherapy by computationally aided real-time personalization. Cancer Res. 2012;72: 2218–2227.

12. Kuznetsov VA, Knott GD. Modeling tumor regrowth and immunotherapy. Math Comput Model. 2001;33: 1275–1287.

13. dePillis LG, Eladdadi A, Radunskaya AE. Modeling cancer-immune responses to therapy. J Pharmacokinet Pharmacodyn. 2014;41: 461–478.

14. Leder K, Pitter K, LaPlant Q, Hambardzumyan D, Ross BD, Chan TA, et al. Mathematical modeling of PDGF-driven glioblastoma reveals optimized radiation dosing schedules. Cell. 2014;156: 603–616.

15. Michor F, Beal K. Improving Cancer Treatment via Mathematical Modeling: Surmounting the Challenges Is Worth the Effort. Cell. 2015;163: 1059–1063.

16. Gompertz B. XXIV. On the nature of the function expressive of the law of human mortality, and on a new mode of determining the value of life contingencies. In a letter to Francis Baily, Esq. F. R. S. &c. Philosophical Transactions of the Royal Society of London. 1825;115: 513–583.

17. Von Bertalanffy L. Quantitative laws in metabolism and growth. Q Rev Biol. 1957;32: 217–231.

18. Laird AK. DYNAMICS OF TUMOUR GROWTH: COMPARISON OF GROWTH RATES AND EXTRAPOLATION OF GROWTH CURVE TO ONE CELL. Br J Cancer. 1965;19: 278–291.

19. Norton L, Simon R, Brereton HD, Bogden AE. Predicting the course of Gompertzian growth. Nature. 1976;264: 542–545.

20. Benzekry S, Lamont C, Beheshti A, Tracz A, Ebos JML, Hlatky L, et al. Classical mathematical models for description and prediction of experimental tumor growth. PLoS Comput Biol. 2014;10: e1003800.

21. Mehrara E, Forssell-Aronsson E, Ahlman H, Bernhardt A. Specific Growth Rate versus Doubling Time for Quantitative Characterization of Tumor Growth Rate. Cancer Res. 2007;67: 3970–3975.

22. Vaidya VG, Alexandro FJ Jr. Evaluation of some mathematical models for tumor growth. Int J Biomed Comput. 1982;13: 19–36.

23. Hutter C, Zenklusen JC. The Cancer Genome Atlas: Creating Lasting Value beyond Its Data. Cell. 2018;173: 283–285.

24. Taylor AM, Shih J, Ha G, Gao GF, Zhang X, Berger AC, et al. Genomic and Functional Approaches to Understanding Cancer Aneuploidy. Cancer Cell. 2018;33: 676–689.e3.

25. Thorsson V, Gibbs DL, Brown SD, Wolf D, Bortone DS, Ou Yang T-H, et al. The Immune Landscape of Cancer. Immunity. 2018;48: 812–830.e14.

26. Berger AC, Korkut A, Kanchi RS, Hegde AM, Lenoir W, Liu W, et al. A Comprehensive Pan-Cancer Molecular Study of Gynecologic and Breast Cancers. Cancer Cell. 2018;33: 690–705.e9.

27. Vaghi C, Rodallec A, Fanciullino R, Ciccolini J, Mochel JP, Mastri M, et al. Population modeling of tumor growth curves and the reduced Gompertz model improve prediction of the age of experimental tumors. PLoS Comput Biol. 2020;16: e1007178.

28. Mak IW, Evaniew N, Ghert M. Lost in translation: animal models and clinical trials in cancer treatment. Am J Transl Res. 2014;6: 114–118.

29. Ruggeri BA, Camp F, Miknyoczki S. Animal models of disease: pre-clinical animal models of cancer and their applications and utility in drug discovery. Biochem Pharmacol. 2014;87: 150–161.

30. Brady R, Enderling H. Mathematical Models of Cancer: When to Predict Novel Therapies, and When Not to. Bull Math Biol. 2019;81: 3722–3731.

31. Falzone L, Salomone S, Libra M. Evolution of Cancer Pharmacological Treatments at the Turn of the Third Millennium. Front Pharmacol. 2018;9: 1300.

32. Collins GS, Reitsma JB, Altman DG, Moons KGM. Transparent Reporting of a multivariable prediction model for Individual Prognosis or Diagnosis (TRIPOD): the TRIPOD statement. Ann Intern Med. 2015;162: 55–63.

33. Faustino-Rocha A, Oliveira PA, Pinho-Oliveira J, Teixeira-Guedes C, Soares-Maia R, da Costa RG, et al. Estimation of rat mammary tumor volume using caliper and ultrasonography measurements. Lab Anim. 2013;42: 217–224.

34. Therasse P, Arbuck SG, Eisenhauer EA, Wanders J, Kaplan RS, Rubinstein L, et al. New guidelines to evaluate the response to treatment in solid tumors. European Organization for Research and Treatment of Cancer, National Cancer Institute of the United States, National Cancer Institute of Canada. J Natl Cancer Inst. 2000;92: 205–216.

35. Eisenhauer EA, Therasse P, Bogaerts J, Schwartz LH, Sargent D, Ford R, et al. New response evaluation criteria in solid tumours: revised RECIST guideline (version 1.1). Eur J Cancer. 2009;45: 228–247.

36. Fleetwood K. An introduction to differential evolution. Proceedings of Mathematics and Statistics of Complex Systems (MASCOS) One Day Symposium, 26th November, Brisbane, Australia. maths.uq.edu.au; 2004. pp. 785–791.

37. Bagchi S, Yuan R, Engleman EG. Immune Checkpoint Inhibitors for the Treatment of Cancer: Clinical Impact and Mechanisms of Response and Resistance. Annu Rev Pathol. 2021;16: 223–249.

38. Ferrara R, Pilotto S, Caccese M, Grizzi G, Sperduti I, Giannarelli D, et al. Do immune checkpoint inhibitors need new studies methodology? J Thorac Dis. 2018;10: S1564–S1580.

39. Soria F, Beleni AI, D’Andrea D, Resch I, Gust KM, Gontero P, et al. Pseudoprogression and hyperprogression during immune checkpoint inhibitor therapy for urothelial and kidney cancer. World J Urol. 2018;36: 1703–1709.

40. Kataoka Y, Hirano K. Which criteria should we use to evaluate the efficacy of immune-checkpoint inhibitors? Annals of translational medicine. 2018. p. 222.

41. Kuang Y, Nagy JD, Eikenberry SE. Introduction to Mathematical Oncology. CRC Press; 2018.

42. Noble R, Burri D, Kather JN, Beerenwinkel N. Spatial structure governs the mode of tumour evolution. bioRxiv. 2019. Available: https://www.biorxiv.org/content/10.1101/586735v1.abstract

43. Jarrett AM, Lima EABF, Hormuth DA 2nd, McKenna MT, Feng X, Ekrut DA, et al. Mathematical models of tumor cell proliferation: A review of the literature. Expert Rev Anticancer Ther. 2018;18: 1271–1286.

44. Collin A, Copol C, Pianet V, Colin T, Engelhardt J, Kantor G, et al. Spatial mechanistic modeling for prediction of the growth of asymptomatic meningiomas. Comput Methods Programs Biomed. 2021;199: 105829.

45. Kather JN, Poleszczuk J, Suarez-Carmona M, Krisam J, Charoentong P, Valous NA, et al. In Silico Modeling of Immunotherapy and Stroma-Targeting Therapies in Human Colorectal Cancer. Cancer Res. 2017;77: 6442–6452.

46. Kather JN, Charoentong P, Suarez-Carmona M, Herpel E, Klupp F, Ulrich A, et al. High-Throughput Screening of Combinatorial Immunotherapies with Patient-Specific In Silico Models of Metastatic Colorectal Cancer. Cancer Res. 2018;78: 5155–5163.

47. You L, Brown JS, Thuijsman F, Cunningham JJ, Gatenby RA, Zhang J, et al. Spatial vs. non-spatial eco-evolutionary dynamics in a tumor growth model. J Theor Biol. 2017;435: 78–97.

48. West J, Robertson-Tessi M, Luddy K, Park DS, Williamson DFK, Harmon C, et al. The Immune Checkpoint Kick Start: Optimization of Neoadjuvant Combination Therapy Using Game Theory. JCO Clin Cancer Inform. 2019;3: 1–12.

49. Bayer P, Brown JS, Stańková K. A two-phenotype model of immune evasion by cancer cells. J Theor Biol. 2018;455: 191–204.

50. Stanková K, Brown JS, Dalton WS, Gatenby RA. Optimizing Cancer Treatment Using Game Theory: A Review. JAMA Oncol. 2019;5: 96–103.

51. Gerlee P. The model muddle: in search of tumor growth laws. Cancer Res. 2013;73: 2407–2411.

52. Spigel DR, Chaft JE, Gettinger S, Chao BH, Dirix L, Schmid P, et al. FIR: Efficacy, Safety, and Biomarker Analysis of a Phase II Open-Label Study of Atezolizumab in PD-L1-Selected Patients With NSCLC. J Thorac Oncol. 2018;13: 1733–1742.

53. Fehrenbacher L, Spira A, Ballinger M, Kowanetz M, Vansteenkiste J, Mazieres J, et al. Atezolizumab versus docetaxel for patients with previously treated non-small-cell lung cancer (POPLAR): a multicentre, open-label, phase 2 randomised controlled trial. Lancet. 2016;387: 1837–1846.

54. Peters S, Gettinger S, Johnson ML, Jänne PA, Garassino MC, Christoph D, et al. Phase II Trial of Atezolizumab As First-Line or Subsequent Therapy for Patients With Programmed Death-Ligand 1-Selected Advanced Non-Small-Cell Lung Cancer (BIRCH). J Clin Oncol. 2017;35: 2781–2789.

55. Rittmeyer A, Barlesi F, Waterkamp D, Park K, Ciardiello F, von Pawel J, et al. Atezolizumab versus docetaxel in patients with previously treated non-small-cell lung cancer (OAK): a phase 3, open-label, multicentre randomised controlled trial. Lancet. 2017;389: 255–265.

56. Balar AV, Galsky MD, Rosenberg JE, Powles T, Petrylak DP, Bellmunt J, et al. Atezolizumab as first-line treatment in cisplatin-ineligible patients with locally advanced and metastatic urothelial carcinoma: a single-arm, multicentre, phase 2 trial. Lancet. 2017;389: 67–76.

57. Gyllenberg M, Webb GF. Quiescence as an explanation of Gompertzian tumor growth. Growth Dev Aging. 1989;53: 25–33.

