## Supplementary figures and images for "Classical Mathematical Models for Prediction of Response to Chemotherapy and Immunotherapy"

### Suppl. Figure 1

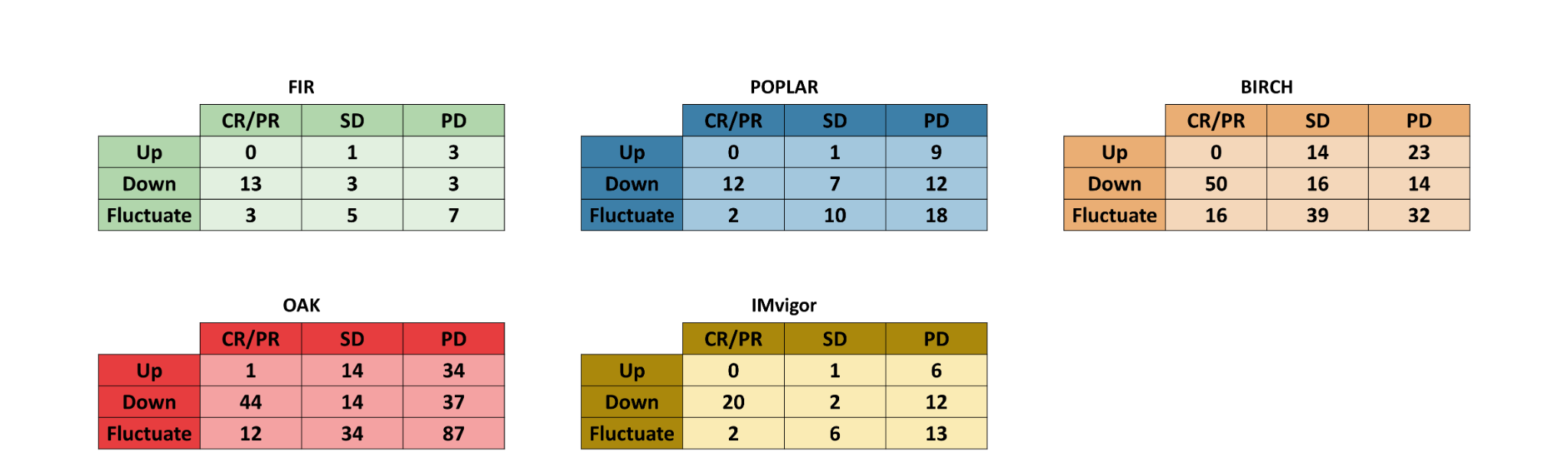

### Suppl. Figure 2

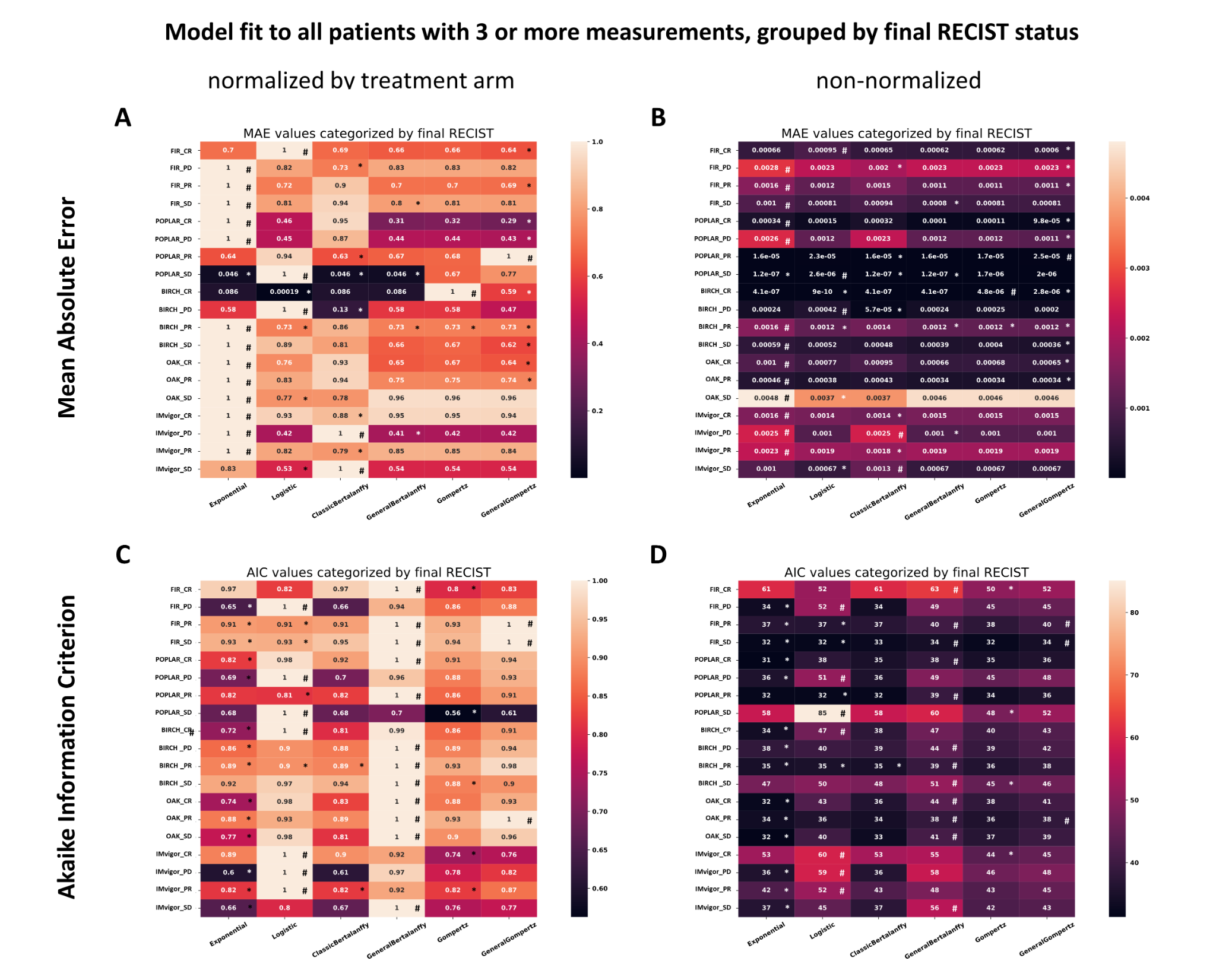

### Suppl. Figure 3

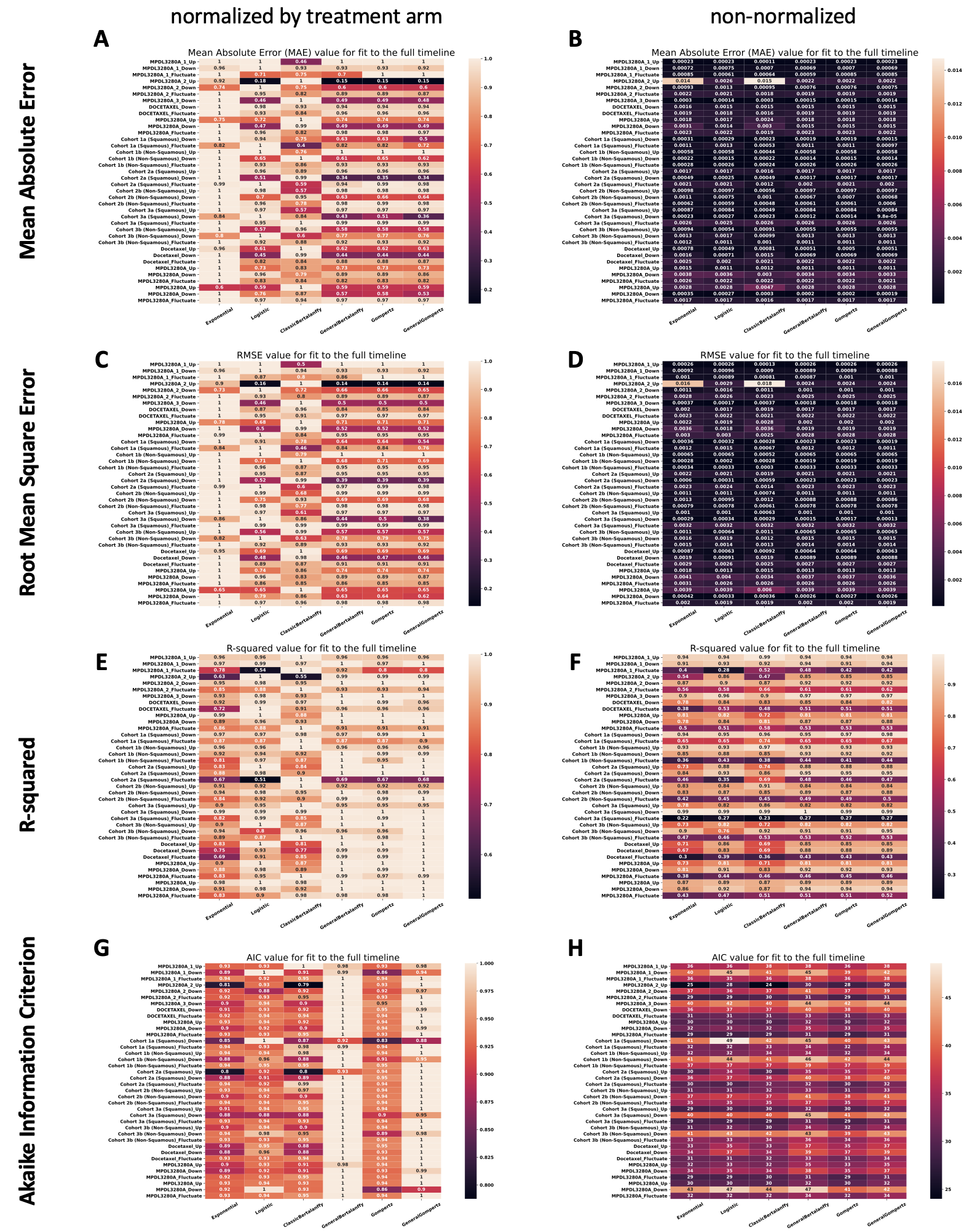

### Suppl. Figure 4

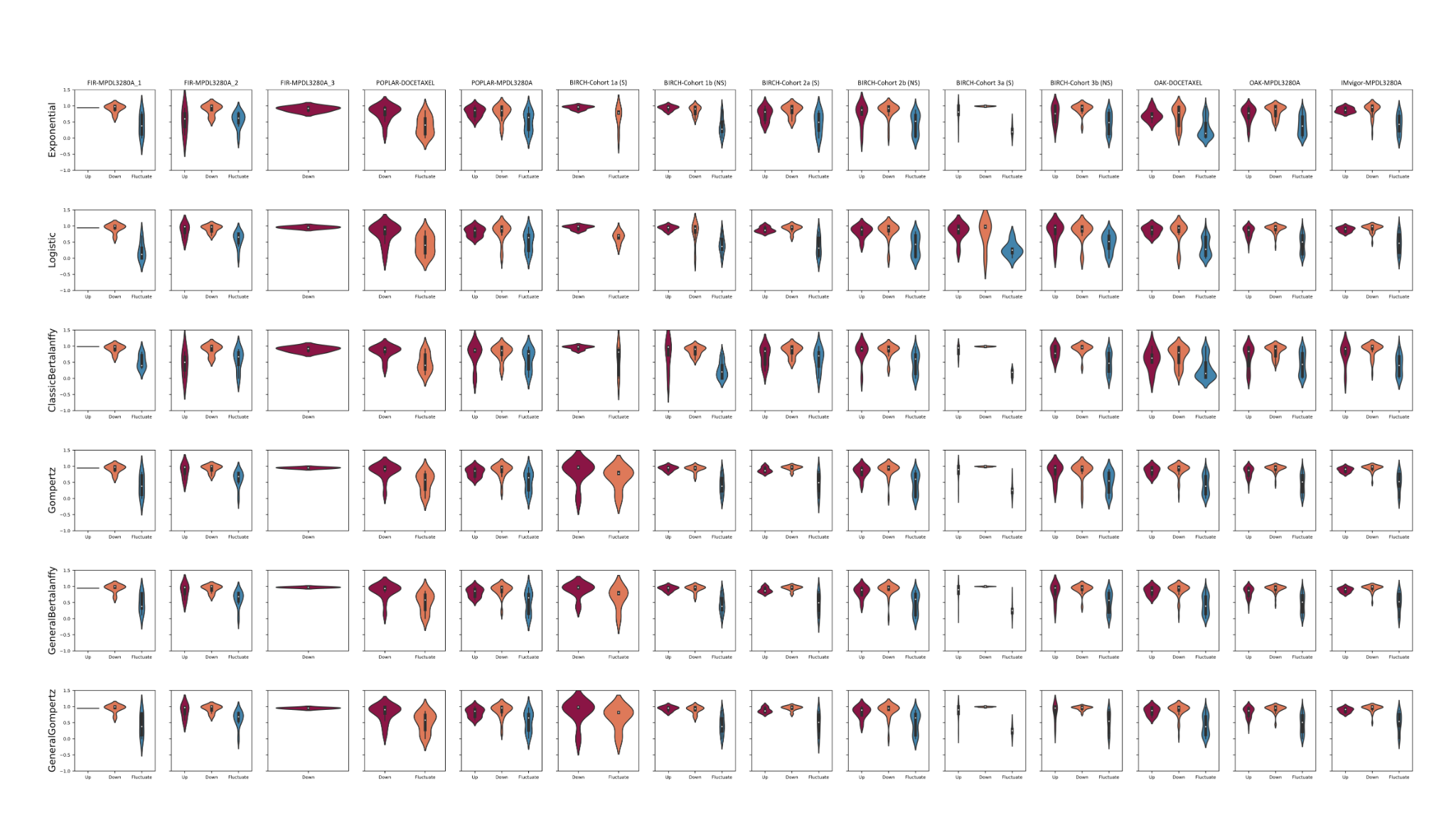

### Suppl. Figure 5

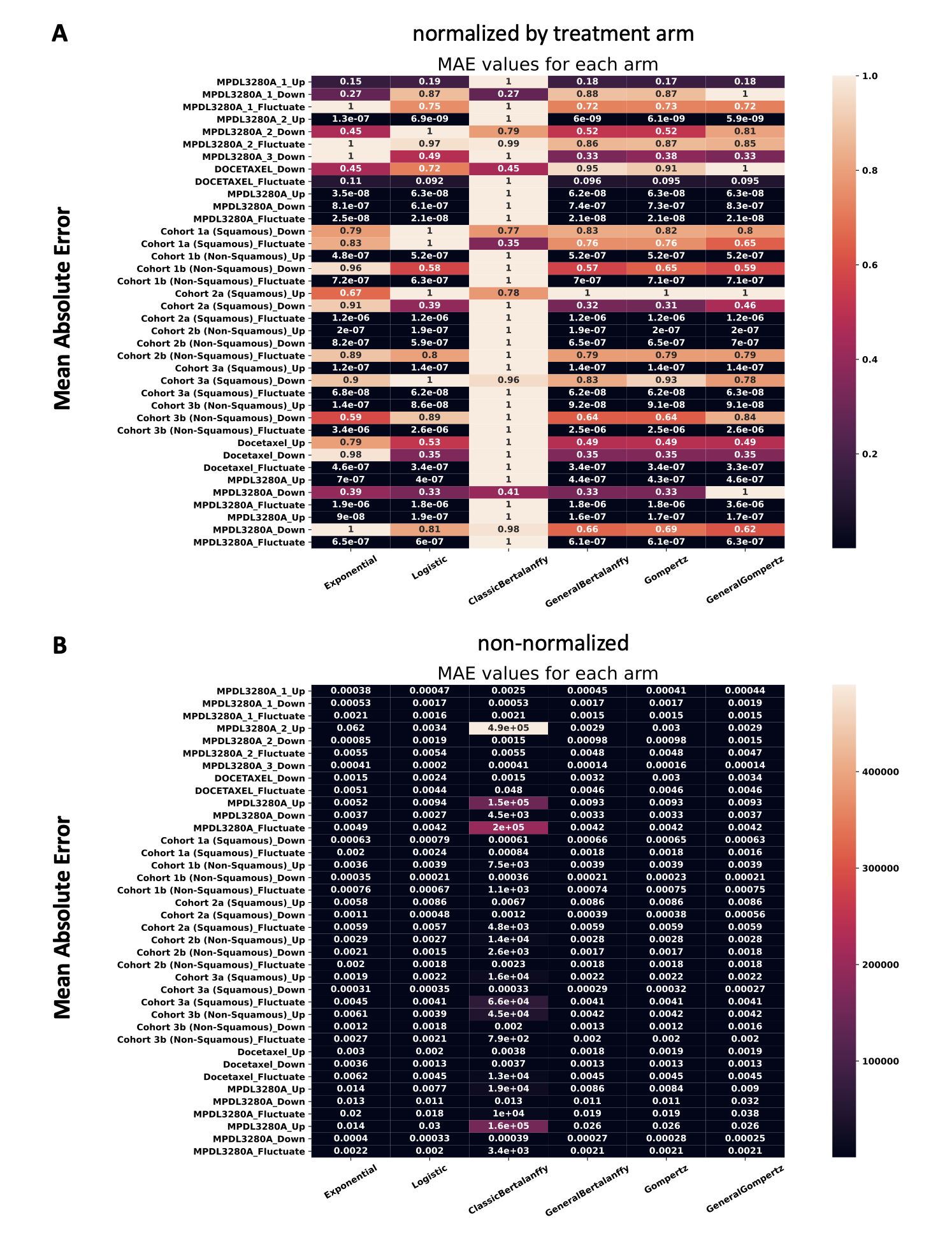
